# Travelling with a parasite: the evolution of resistance and dispersal syndromes during experimental range expansion

**DOI:** 10.1101/2020.01.29.924498

**Authors:** Giacomo Zilio, Louise S. Nørgaard, Claire Gougat-Barbera, Matthew D. Hall, Emanuel A. Fronhofer, Oliver Kaltz

**Affiliations:** ISEM, University of Montpellier, CNRS, EPHE, IRD, Montpellier, France; School of Biological Sciences and Centre for Geometric Biology, Monash University, Melbourne, Australia

**Keywords:** Dispersal syndromes, eco-evolutionary dynamics, host-parasite interactions, invasion front, range expansion, resistance

## Abstract

Rapid evolutionary change during range expansions can lead to diverging range core and front populations, with the emergence of dispersal syndromes (coupled responses in dispersal and life-history traits). Besides intraspecific effects, range expansions may be impacted by interspecific interactions such as parasitism. Yet, despite the potentially large impact of parasites imposing additional selective pressures on the host, their role on range expansions remains largely unexplored. Using microcosm populations of the ciliate *Paramecium caudatum* and its bacterial parasite *Holospora undulata*, we studied experimental range expansions under parasite presence or absence. We found that the interaction of range expansion and parasite treatments affected the evolution of host dispersal syndromes. Namely, front populations showed different associations of population growth parameters and swimming behaviours than core populations, indicating divergent evolution. Parasitism reshaped trait associations, with hosts evolved in the presence of the parasite exhibiting overall increased resistance and reduced dispersal. Nonetheless, when comparing infected range core and front populations, we found a positive association, suggesting joint evolution of resistance and dispersal at the front. We conclude that host-parasite interactions during range expansions can change evolutionary trajectories; this in turn may feed back on the ecological dynamics of the range expansion and parasite epidemics.

## Introduction

Species range expansions are increasing in frequency in response to rapidly changing environments [1] and becoming crucial to reducing the risk of extinction [2]. While ecological and evolutionary processes have been well explored for single species, we are still lacking a good community perspective of range shifts [3,4]. Species might rarely “travel alone”, and it is conceivable that interactions with other organisms facilitate or slow down spatial spread, and add new selection pressures while a species is spreading.

Antagonistic species interactions have been suggested to be a major factor defining species ranges [5,6]. Parasitism, for example, are ubiquitous and exert strong demographic and evolutionary pressures on the host [7,8], which can limit geographic ranges as has been shown in some theoretical models [9–11]. Parasites may re-enforce range limits by reducing dispersal and/or by imposing additional mortality in already small populations at the range front. Such modifications of the ecological dynamics seems to limit the spread of the invasive cane toad (*Rhinella marina*) in Australia [12,13], and also play a role in other natural systems [14–16]. The evolution of counter-adaptation against parasites may remove such range limits, but also lead to more complex outcomes if parasite-mediated selection affects the evolution of other life-history traits known to be selected at range fronts. Novel trait combinations may then produce eco-evolutionary feedbacks influencing rates of spatial spread of both the host species and the parasite. This may cause knock-on effects on interactions with other species in the community [17,18] or increase the risk of spill-over of disease [19,20]. Thus, understanding how parasites shape evolutionary trajectories of phenotypic traits in a range expansion context is also of major interest to conservation, disease control [21–23] or invasive species management [24].

Range expansions often involve stochastic genetic change in small vanguard front populations (via drift or founder bias), but also the concerted evolution of dispersal, life-history, behaviour or physiology, often referred to as “dispersal syndrome” [25,26]. Typically range front populations may be populated by individuals with a coloniser phenotype, characterised, e.g., by high dispersal propensity and high reproductive rate. Kin selection and spatial assortative mating are then expected to reinforce such trait associations while the range front is expanding [27,28]. There is empirical and experimental evidence that such spatial selection is indeed a strong evolutionary force. Dispersal is a heritable trait in various biological systems and covariation with other traits has a genetic basis [29], making it possible for selection to drive multi-trait divergence between core and front populations [30,31]. An example is the above-mentioned cane toad, where the evolution of dispersal and other life-history traits promoted the speed of spread at the invasion front [32–34]. Similar results were found for the invasive ladybird *Harmonia axyridis* [35], and in laboratory range expansions using experimental evolution approaches [36–38].

We currently know relatively little about parasite-mediated selection at range fronts and its consequences for dispersal syndrome evolution [3,4]. While a considerable bulk of literature exists on resistance evolution in host-parasite metapopulations [39,40], there is no specific theory in the context of range expansion. Intuitively, we may expect that virulent parasites select for increased resistance and that this removes a parasite-imposed range speed limit. Such a case of resistance evolution has been found for populations of the cane toad infested by lung-worms at the range margin [41,42]. Theory further suggests that natural enemies select for changes in host dispersal. For example, parasite-induced fluctuations in host population dynamics can modify environmental predictability and fitness expectations, and consequently favour increased host dispersal [43,44]. Parasites may also select for plastic responses in dispersal, with sometimes counter-intuitive consequences for the spread of infection [45].

However, few if any studies have considered the joint action of parasite-mediated selection and spatial selection, and hence the interplay between the evolution of resistance and dispersal-related traits. Indeed, selection for resistance can be rapid and strong, but also modify genetic correlations with other life-history traits [46,47], often involving trade-offs [48,49] that constrain evolutionary trajectories. Hence, if selection for resistance is uncorrelated with dispersal syndrome traits (or even positively correlated), counter-adaptation against the natural enemy may simply remove the brake from range expansion. However, if resistance trades off with expansion-relevant traits, such as dispersal or growth, the additional parasite-mediated selection may shift populations to novel positions in multi-trait space, which finally may or may not impede spatial spread.

Using experimental evolution, we investigated the effect of a parasite on the emergence of dispersal syndromes. In a long-term experiment, we mimicked range expansions in interconnected microcosms of the freshwater protist *Paramecium caudatum* and its bacterial parasite *Holospora undulata*. In a range front treatment, the paramecia were constantly selected to disperse into a new microcosm, whereas in the range core treatment, only the non-dispersing fraction of the population was maintained. Range and core treatments were established for infected and uninfected populations. Evolved hosts were assayed for six traits, including dispersal, resistance, population growth characteristics and swimming behaviour. We found substantial divergence in multiple phenotypes in the trait space, attributable to the combined effect of our experimental treatments. A dispersal syndrome emerged, with higher population equilibrium density and movement tortuosity in the front treatment, and higher population growth rate and swimming speed in the core treatment. Evolution with the parasite affected some of these trait relationships and additionally left signatures in the observed levels of resistance and dispersal. Populations evolving with the parasite were more resistant, in particular those in the range front treatment. In contrast, *Paramecium* originating from parasitised populations generally dispersed less than their parasite-free counterparts. The parasite-driven evolution of such novel multi-trait associations may influence the speed of range expansions, but also the risk of spreading epidemics, which can have important implications for biological control or conservation management.

## Material and methods

### Study system

*Paramecium caudatum* is a freshwater filter-feeding ciliate with a world-wide distribution [50]. Ciliates typically exhibit a nuclear dimorphism: The “germ-line” micronucleus is active during the sexual stage, while the highly polyploid “somatic” macronucleus regulates gene expression during the asexual stage, when replication occurs through mitotic division. In this experiment, clonal populations were maintained asexually (max. 1-2 population doublings per day at constant 23°C) in 50 mL tubes, using a sterilised lettuce medium (1g dry weight of organic lettuce per 1.5l of Volvic™ mineral water), supplemented *ad libitum* with the bacterium *Serratia marcescens* as a food resource (referred to as “bacterised medium” or “medium”, hereafter). The gram-negative bacterium *Holospora undulata* is an obligate parasite, infecting the micronucleus of *P. caudatum* [51]. The infection life cycle comprises both horizontal and vertical transmission. *Paramecium* ingest infectious forms from the aquatic environment, which subsequently colonise the micronucleus and differentiate into multiplying reproductive forms; these reproductive forms are vertically transmitted to the daughter cells of mitotically dividing hosts. The infection life cycle is completed when reproductive forms differentiate into infectious forms (c. 7 days post infection), which are then released during host cell division or upon host death. With accumulating parasite loads, infection reduces cell division and survival of the *Paramecium*, as well as dispersal [52–54]. Experimental evolution of resistance to this parasite was demonstrated in previous long-term experiments and can come at reproductive costs [55,56].

### Experimental protocols

#### (i) Long-term range expansion experiment

##### Dispersal in two-patch systems

We used two-patch systems for this selection experiment (Supplementary Information, Fig. S1). The systems were built from two 14 mL plastic tubes (“core patch” and “front patch”) interconnected by 5-cm silicon tubing (0.6 mm inner diameter) serving as a corridor through which the *Paramecium* can actively swim. We define dispersal as the active displacement of *P. caudatum* from the core patch to the front patch. In the long-term experiment, short episodes of dispersal (3h) alternated with periods of population growth and maintenance (1 week). For a dispersal episode, we filled the two-patch system with 9.5 mL of fresh bacterised medium and then blocked the corridor with a clamp. The core patch was filled with c. 8ml of culture containing *Paramecium* (≈ 2000 individuals), whereas the front patch was only topped up with non-bacterised medium and was thus “empty”. After removal of the clamp, *Paramecium* could freely disperse to the front patch or to stay in the core. After three hours, we blocked the corridor and estimated the cell density in the core and front patch, by sampling up to 1mL from each tube and counting the number of individuals under a dissecting microscope. The dispersal rate is thus the number of dispersers divided by the total number of individuals in the two-patch systems, divided by 3 hours.

##### Range expansion treatments

Two “range expansion” treatments were imposed. In the range front treatment, only *Paramecium* that dispersed into the front patch were maintained and allowed to grow for 1 week until the next episode of dispersal. Conversely, in the range core treatment, only the non-dispersing *Paramecium* were maintained and allowed to regrow. These contrasting selection protocols were continued for a total of 26 cycles. The front selection treatment mimics the leading front of range expansion or a biological invasion, with populations continuously dispersing into a new microcosm. Populations from the range core treatment stay in place and continuously lose emigrants. Each new growth cycle was started by placing on average 200 paramecia from front and core treatments in 20 mL of fresh bacterised medium, and equilibrium density was then reached within the following 3-4 days. The experiment was conducted with a single host line [53]. This line (63D) had undergone three years of parasite-free core selection prior to the present experiment; initially started from a mix of strains, it has become fixed for a single cytochrome oxydase I haplotype [57].

##### Parasite treatment

Range core and front treatments were established for both infected and uninfected populations. The parasites were taken from an experiment [53] that had already been imposing core and front selection on infected 63D populations for about 8 months (30 cycles). Using standard protocols (see S2), we extracted infectious forms of the parasite from 5 core selection lines and from 5 front selection lines, which were then used to inoculate our new, naive 63D hosts. In other words, we continued range core and front treatments for the parasite, but replaced the previous hosts by new unselected hosts. In addition to these 10 infected selection lines, we established 3 uninfected front-selection lines and 3 uninfected core-selection lines as controls, giving a total of 16 selection lines. After this initial inoculation of the experimental lines, we did not interfere with epidemiological dynamics and all transmission occurred naturally over the course of the 26 new cycles of the long-term experiment. We routinely measured dispersal and population density (Fig. S4) and verified the presence of infection in the parasite treatment.

#### (ii) Phenotypic trait assays

At the end of the long-term experiment, phenotypic trait assays for *Paramecium* from all 16 selection lines were performed under common-garden conditions. Using a micropipette, we arbitrarily picked 4 uninfected paramecia from each selection line and placed them individually in single 1.5 mL Eppendorf tubes filled with bacterised medium, where they were allowed to grow for 2 weeks until small monoclonal lines had established (c. 7-8 asexual generations). Each monoclonal line was then split into three technical replicates and grown for a second common-garden period of 10 days in 50-mL tubes to obtain mass cultures for the phenotype assays (16 selection lines x 4 monoclonal lines x 3 technical replicates = 64 monoclonal lines and 192 replicates, Fig. S3). Of the 64 monoclonal lines, one monoclonal line from the core treatment with parasite did not grow and was lost, leaving us with 63 lines available to measure 6 phenotypic traits, as follows.

##### Resistance

To measure resistance, the *Paramecium* were confronted with parasites from range core and front treatments. To this end, we prepared inocula by extracting infectious forms from mixes of the 5 infected core and the 5 infected front selection lines (for details of the protocol, see S2). For inoculation, c. 5000 paramecia were placed in a volume of 25 mL in a 50-mL tube, to which we added 4.5 × 10^5^ infectious forms (core-parasite or front-parasite inoculum). In this way, we set up 4-8 inoculated tubes per host selection line, balanced between the two parasite inocula (16 selection lines x 2-4 monoclonal lines x 2 technical replicates = 120 inoculated tubes). Four days post-inoculation, we fixed c. 20 individuals from each inoculated replicate with lacto-aceto-orcein [51] and inspected them for absence or presence of infection using a phase-contrast microscope (1000x magnification). We define resistance as the proportion of uninfected individuals in the sample. Preliminary analysis showed that *Paramecium* from the four different selection treatments did not differ in their resistance to the mixes of front or core parasites (F_3,12_ = 0.44, n.s.); we therefore combined the two inoculum sources into a single “infected” category for the main analysis. Note, however, that this does not rule out the possibility of specific responses to range or core parasites in other host traits.

##### Dispersal rate

Dispersal was measured in linear 3-patch systems (50 mL tubes; Fig. S5), where uninfected *Paramecium* dispersed from the middle tube into the two outer tubes (see SI for detailed protocol). This system configuration allowed us to use bigger volumes of culture and thus obtain higher numbers of dispersers than in 2-patch systems. Connections were opened for 3 h, dispersal rates were then estimated by counting the *Paramecium* in samples from the central tube (500 μl) and from the combined two outer tubes (3 mL). We employed technical replicates that had not been used for the resistance assay, and were kept in 30 mL of fresh medium for several days prior to the dispersal test. One replicate per monoclonal line was tested (63 dispersal tests).

##### Population growth rate and equilibrium density

For the population growth assay, we placed groups of 5 arbitrarily picked *Paramecium* in 15-mL tubes filled with 10 mL of bacterised medium. Over 9 days, we tracked densities in 24-h intervals, estimated from the number of individuals present in 200-μL samples. We set up 6-12 tubes per host selection line (3 technical replicates per monoclonal line), with a total of 180 replicates. Only uninfected *Paramecium* were tested. For each tube, estimates of intrinsic population growth rate (*r*_*0*_) were obtained by fitting a Beverton-Holt population growth model to each density time series, using a Bayesian approach [58],. For certain tubes we obtained unsatisfactory fits of equilibrium density; we therefore decided to use the mean density over the second half of the assay (day 5-9) as a proxy for equilibrium density 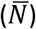. A small fraction of 19 tubes (corresponding to 9 monoclonal lines from selection lines from range front and parasitism treatment) failed to grow and remained at very low density for unknown reasons; it was not possible to fit our population growth model to these data, and these replicates were therefore excluded from analysis.

##### Swimming speed and tortuosity

At the end of the above population growth assay, we assessed swimming behaviour, using an established pipeline of computer vision and automated video analysis to collect this data [59,60]. From a given tube, one sample of 119 μL was imaged under a Perfex Pro 10 stereomicroscope, using a Perfex SC38800 camera (15 frames per second; duration: 10 s; total magnification: 10x). Videos were analysed using the bemovi R-package [60], which provided individual-based data on swimming speed and the tortuosity of swimming trajectories (standard deviation of the turning angle distribution). Swimming speed and tortuosity were averaged over all individuals in a sample prior to analysis. A total of 63 samples (1 per monoclonal line) was used for analysis, giving 2-4 observations per host selection line. Three monoclonal lines (front parasite-exposed) went extinct prior to video recording.

### Statistical analysis

All statistical analyses were performed with R v 4.2.0 [61], using Bayesian models with the “rstan” (version 2.21.5), “brms” (version 2.17.0) and “rethinking” (version 2.21) packages [62–64]. Focusing on the analysis of trait associations, we constructed a data matrix with the measurements of the 6 traits for 63 monoclonal lines (for resistance, the mean over the two technical replicates was calculated). To impute 24 missing observations (9 for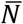and *r*_*0*_, 3 for swimming speed and tortuosity), we used the “mice” package version 3.14.0 [65] and took the mean of 1000 multiple imputations (24/378 imputed values). Trait distributions were standardized by using the standardize function of the “rethinking” package [64].

We then fitted five multivariate multilevel models, from the intercept to the full interaction model, using a normal error structure (chain length: warmup = 1,000 iterations, chain = 20,000 iterations). The explanatory factors of the full model were range expansion treatment (core vs front selection lines), parasite treatment (infected vs uninfected control selection lines) and their interaction, selection line identity was included as random term. Models were fitted with the default vaguely informative priors of the “brms” package. We compared and ranked the five models using the Watanabe–Akaike information criterion, WAIC [66], a generalized version of the Akaike information criterion [67]. We used a conservative approach based on WAIC weights and not on the best ranked model. We averaged the posterior predictions of the models and calculated the relative importance (RI) of the explanatory variables. The RI corresponds to the sum of the respective WAIC model weights in which the explanatory variable was present. From this same data set and following the same procedure we also performed univariate analyses for each of the 6 traits. In addition, for a complementary graphical inspection of the results we used the above data matrix to carry out a principal component analysis (PCA).

## Results

Multivariate analysis revealed signatures of selection history, with strong effects of range expansion treatment (relative importance, RI = 1.00) and parasitism treatment (RI = 0.77) on the observed phenotypic variation and covariation (Table 1). After model selection, the interaction between range expansion and parasitism (RI = 0.55) was retained in the best model fit (lowest WAIC; Model 5 in Table 1), indicating that the effect of spatial selection acted jointly with the presence of the parasite. In additional univariate analyses, all 6 traits showed signals of the two experimental treatments in combination or alone (Table 2, details for all traits in SI, Table S3-S8).

**Table 1.** Statistical models and parameters included in the main multivariate trait analysis and model averaging. Rows are the different models; the best model is highlighted in bold. Columns are the explanatory variables included with the corresponding WAIC, standard error of the WAIC and WAIC weights for each model. The RI row indicates the relative importance of the explanatory variables.

|  | Range expansion | Parasitism | Range * Parasitism | WAIC | SE | WAIC weight |
| --- | --- | --- | --- | --- | --- | --- |
| Model 1 |  |  |  | 951.2 | 40.2 | 0.00 |
| Model 2 |  | X |  | 952.3 | 39.5 | 0.00 |
| Model 3 | X |  |  | 942.1 | 40.0 | 0.23 |
| Model 4 | X | X |  | 942.2 | 39.5 | 0.22 |
| **Model 5** | **X** | **X** | **X** | **940.4** | **39.1** | **0.55** |
| RI | 1.00 | 0.77 | 0.55 |  |  |  |

**Table 2.** Best model and WAIC weights from model averaging of single trait analysis. The explanatory variables included in the best model are indicated with X. The complete list of the models for each trait is given in the SI.

|  | Range expansion | Parasitism | Range * Parasitism | WAIC weight |
| --- | --- | --- | --- | --- |
| Dispersal |  | X |  | 0.28 |
| Resistance | X | X | X | 0.42 |
| $r_0$ | X | X | X | 0.26 |
| $\bar{N}$ | X | X | | 0.23 |
| Speed | X |  |  | 0.62 |
| Tortuosity | X |  |  | 0.39 |

PCA visualizes the (co)variation in multivariate trait space (Fig. 1) and the relative contribution of the six measured traits to the observed patterns of divergence among monoclonal lines (individual points in Fig. 1) and combinations of treatments. As shown by the arrows in Fig. 1, the six traits have similar weight but different orientation (Table S2; PCA loadings), with some pointing in opposite directions, indicative of trade-offs. The details of the PCA are provided in Tables S1 and S2.

**Figure 1.**
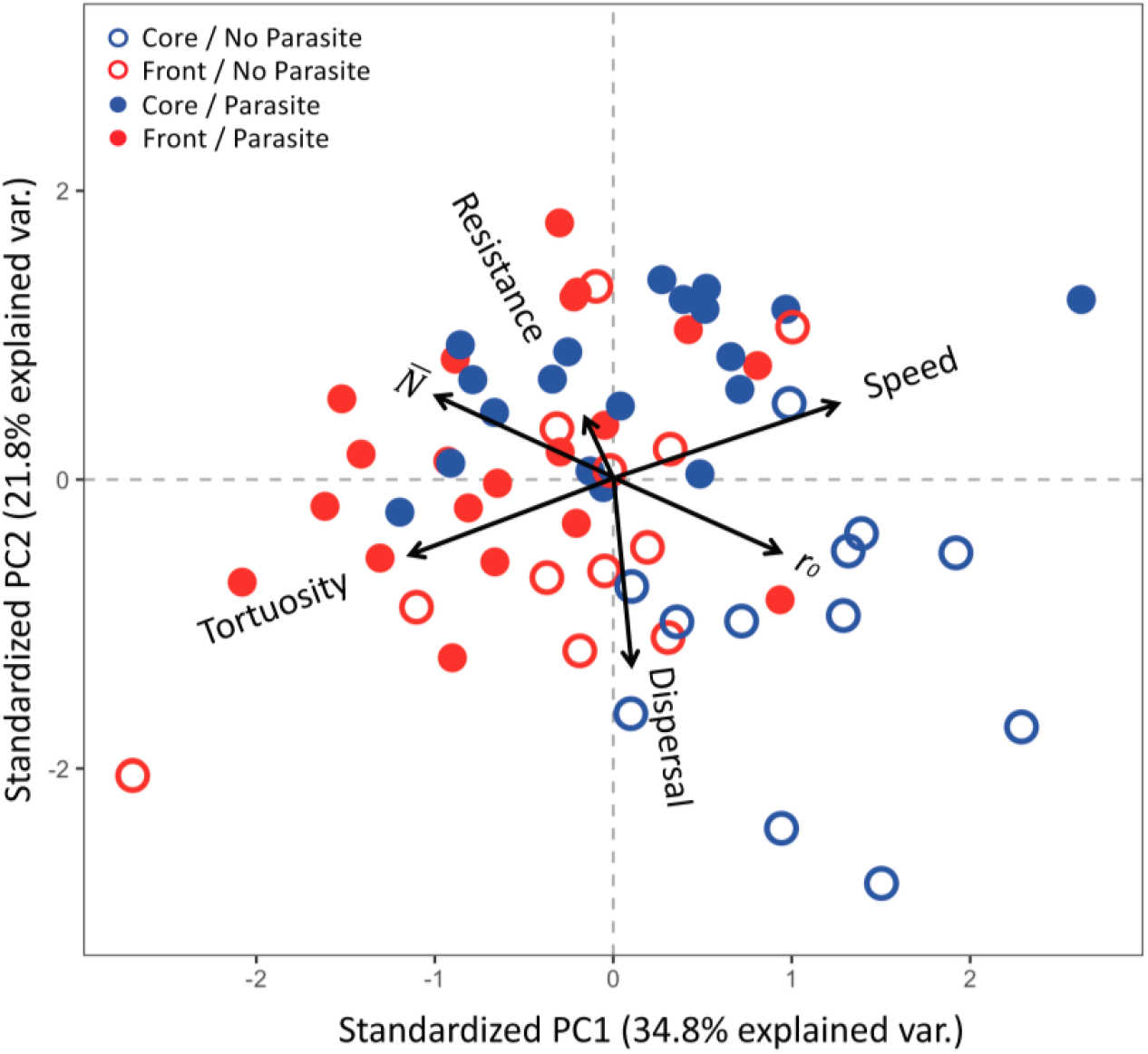
PCA for six traits measured for *P. caudatum* from 4 long-term treatment combinations, showing the first two principal component axes, PC1 and PC2. Each of the 63 points represents the average phenotypic value of a given monoclonal line in multivariate space, with long-term treatment origins specified: range core (blue) vs. range front (red) treatment; evolved in the presence (full circles) vs. absence (empty circles) of the parasite *H. undulata*.

In long-term treatments without parasite, we observed a main pattern of divergence between range core and front treatments along the horizontal PC1 axis (red vs blue open points Fig. 1). The direction and length of the different arrows in Fig. 1 show that this was mainly driven by opposing trends in traits related to demography and movement (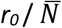and speed / tortuosity have highest PC 1 loadings, Table S2). Accordingly, univariate analyses further revealed high levels of relative importance (RI > 0.58) of the range expansion for these traits (Table S3-S8). Thus, *Paramecium* from the range front populations generally had a 31% lower population growth rate (Fig. 2C) and 54% higher equilibrium density (Fig. 2D) than *Paramecium* from the range core. Similarly, the range front treatment was associated with 12% lower swimming speed (Fig.2E) and 23% more non-linear (i.e., tortuous) movement trajectories (Fig. 2F), compared to the range core treatment.

**Figure 2.**
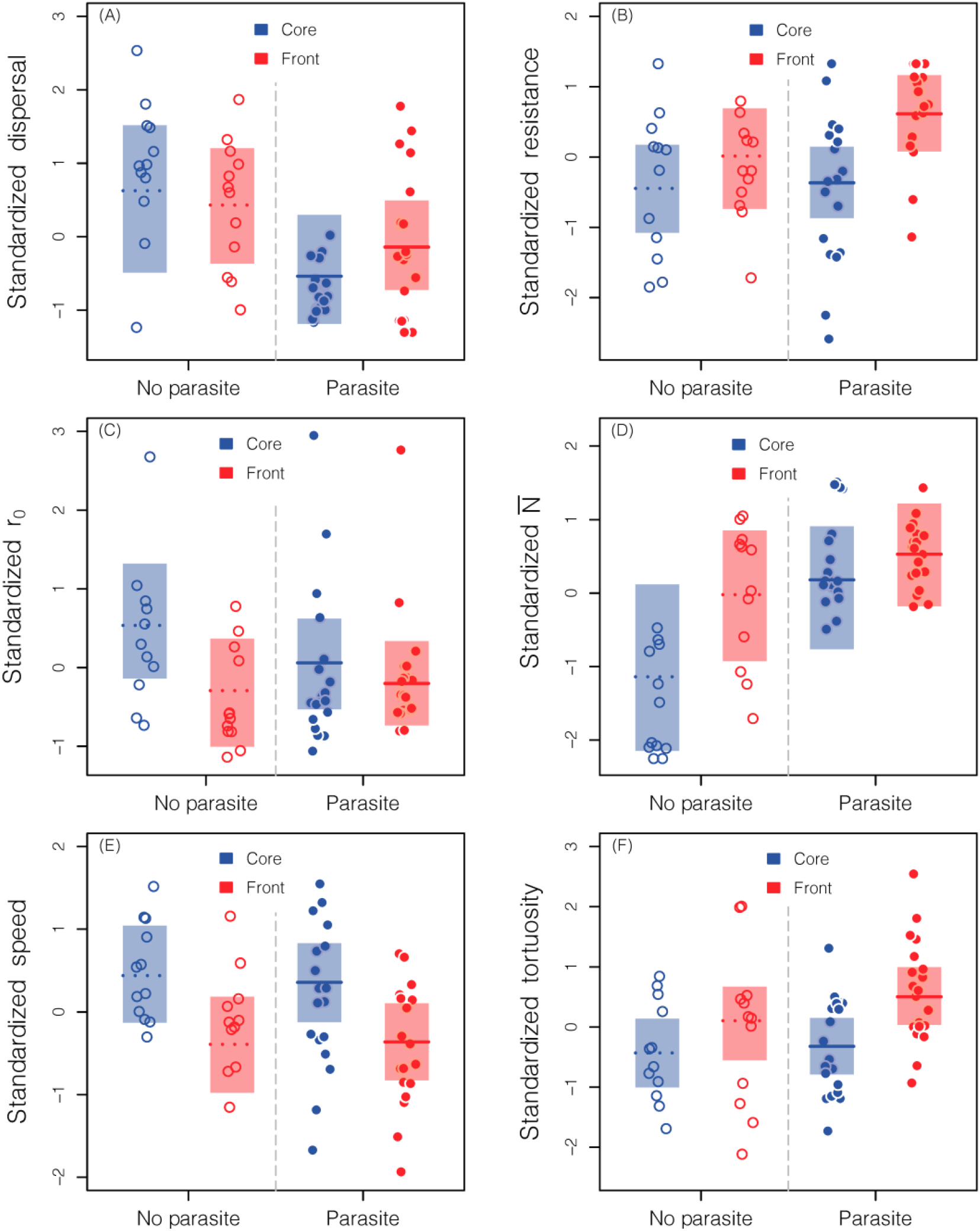
Standardized single trait panels of *P. caudatum* evolved in range core and range front treatments in the presence or absence of parasite: (A) dispersal, (B) resistance, (C) growth rate (*r*_*0*_), (D) equilibrium density 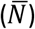, (E) swimming speed and (F) swimming tortuosity. For each panel, red and blue represent core and front treatment while the parasitism treatment is specified as follows: evolved in the presence of the parasite (full dots and lines) vs. parasite-free (empty dots and dashed lines). The points are the means for 63 monoclonal lines, isolated from a total of 16 long-term selection lines (for details, see text). The shaded areas and tick lines are the 95% compatibility interval and median of the averaged model of the posterior distributions.

In long-term treatments with the parasite, phenotypic differentiation between infected and parasite-free lines can be seen along the PC2 axis, and in part along the PC1 axis (Fig. 1, full vs empty dots). To some degree, this parasite effect varied for range and core treatments, leading to overlap of the different treatment combinations in Fig. 1. We identify three patterns. First, resistance and dispersal are the key traits associated with the PC2 axis (Fig. 1), with trait trajectories indicating higher resistance for lines evolved with the parasite and higher dispersal for parasite-free lines. Univariate analyses are in line with these trends. For resistance, there was an interaction between range expansion and parasitism (Table 2 and Table S4; RI interaction = 0.42). Thus, the presence of parasite tended to increase resistance in the range front treatment (+15%), but less so in core treatment (Fig. 2B) compared to the unexposed evolved counterparts (Fig. 2B). For dispersal, there was a main effect of parasite treatment, with a generally lower dispersal of *Paramecium* from infected long-term populations in both range core and front treatments (Table 2 and Table S3; RI parasitism = 0.28; Fig. 2A).

Second, the parasite long-term treatment further modified differences in demographic traits, as shown by the partial separation of parasite and parasite-free treatments along the opposing 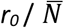 trajectories in the PCA (Fig. 1). Thus, univariate analyses indicate that *Paramecium* from infected populations tended to have higher equilibrium density (Fig. 2D) and lower growth rate (Fig. 2C) than those from parasite-free populations (parasitism main effects, Table S5, S6). We also note that range front vs core differentiation in these traits was less pronounced for infected populations, specifically for *r*_*0*_ (RI range * parasitism interaction = 0.26, Table 2). Third, the long-term presence of the parasite had no obvious effect on swimming behaviour (Fig.2E-F), leaving only the range treatment in the best model in the univariate analysis (RI > 0.93, Table S7, S8).

## Discussion

Eco-evolutionary dynamics in expanding edge populations can favour specific adaptations in dispersal capacity and reproductive strategies (“dispersal syndromes”), but still little is known about how these processes are modulated by interactions with natural enemies or parasites. This study investigated the interplay between spatial selection (range front vs core treatment) and parasite-mediated selection (absence vs presence of parasite) in a factorial experimental design. We found substantial divergence between our experimental range core and front populations, involving traits commonly known to produce “dispersal syndromes” (dispersal, population growth, movement behaviour). In addition, exposure to the parasite in infected populations brought into a play an additional trait (resistance), but also affected divergence in the other traits. In particular, populations evolved with the parasite were more resistant, but dispersed less than their uninfected counterparts. Three pairs of traits acted as opposing vectors in multi-trait space defining the phenotypic divergence between treatments: dispersal - resistance, population growth rate (*r*_*0*_) - equilibrium density 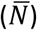 and swimming speed - tortuosity. Below we focus on the evolutionary forces driving these trait associations and discuss possible feedbacks on range expansion speed.

### Resistance and dispersal: trade-off or concerted evolution?

Dispersal and resistance are among the most prominent traits expected to be under selection in range expansion and host-parasite coevolution contexts, respectively. In our study, we investigated their joint evolution. The PCA shows that the two traits have opposite signs, indicating a general negative relationship between dispersal and resistance. This negative association further scales up to the “parasite treatment” level (Fig. 2A, B), suggesting that parasite-mediated selection for increased resistance is accompanied by a decrease in dispersal. This general pattern was most obvious for populations in the range front treatment (panels on the right in Fig. 2A, B), for which we also detected a clear negative quasi-genetic (i.e., across monoclonal lines) correlation between the two traits (Table S9).

Previous work in our study system provided evidence for ample naturally occurring genetic variation in both resistance and dispersal [54,68], and both traits readily evolve under experimental conditions [55,57]. The genetic basis of the two traits is unknown, but we speculate that a trade-off could result from energy constraints arising from mounting an effective (constitutive) defence against the parasite, and/or from concomitant behavioural change. For example, the filter-feeding paramecia become infected when they ingest infectious spores together with other food particles. Hence, contact rates with the parasite may be reduced via reduced feeding activity. Such behavioural aspects of resistance may also include altered swimming behaviour, which can, in turn, affect dispersal (see below).

Even though resistance [69] and dispersal [70] are usually considered costly traits, we do not necessarily always expect trade-offs to evolve. In certain bacteria, cell surface modification conferring resistance to bacteriophages leads to reduced motility [71,72]. Experiments also demonstrated that phage selects for increased bacterial dispersal [71,73]. However, the link between dispersal and resistance was not clear, implying that the two traits evolved independently. This suggests that, apart from mechanical or physiological constraints, the emergence of evolutionary trade-offs also depends on the underlying genetic architecture and relationships with fitness, or on the environmental conditions [45,54,74,75].

Finally, we would like to highlight that trade-offs can be a matter of perspective. A (genetic) trade-off with resistance indeed suggests that parasites constrain the evolution of dispersal syndromes at the range front. Yet, if we compare the two traits between range core and front treatments (rather than between parasite / no parasite treatments), their association is positive: infected front populations have both higher resistance and higher dispersal, when compared with the corresponding core populations (Fig. 2A, B, right panels). In fact, this pattern matches findings for natural populations of the invasive cane toad, where front populations show both higher resistance to lungworm parasites and larger dispersal-related morphological traits [32,41]. From these observations, we would conclude that resistance and dispersal can evolve concomitantly at the invasion front, despite an inherent trade-off between the two traits. It could be the consequence of spatial sorting, putting together variants with generally high levels of resistance and dispersal. In natural systems, similar patterns can be driven by stressful range front environments causing stronger immune responses [76,77].

### Parasitism exacerbates 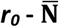 trade-off, but reduces range core-front divergence

On the demographic axis of the multi-trait space, we found opposing directions for equilibrium density 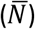 and population growth rate (*r*_*0*_) along the first PC axis, indicating a trade-off typical of classic r-K selection theory (Fig. 1). This axis also separated populations from the range treatment front (high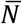, low *r*_*0*_) and range core populations (low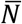, high *r*_*0*_). This pattern seems to contradict common ideas about dispersal syndrome evolution (Burton et al. 2010), predicting competitive strategies in the core (high *K* or 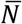) and opportunistic strategies at range fronts (high *r*_*0*_). However, in our system as well as in another protist, competitive ability is characterised by growth rate, rather than equilibrium density [57,59]. Thus, if food bacteria are a slowly regrowing resource, rapacious high-growth protists are favoured in the core treatment and more slowly growing variants at the front [59]. Comparative evidence suggests that this may actually be a widespread pattern [78].

As in the previous section, parasite effects on these demographic traits are a matter of perspective. On the one hand, the 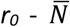 trade-off is more pronounced for populations evolving with the parasite than for parasite-free populations. This trend appears relatively weak for overall treatment means (Fig. 2C, D), but is well supported by supplementary analysis of correlations across monoclonal lines within treatments, showing consistent negative 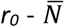 correlations for the parasite treatment (Table S9). Possibly, this pattern reflects indirect responses to selection on resistance, via costs of resistance, acting on demographic traits [79]. On the other hand, the presence of the parasite decreased the trait differences between range core and front treatments, when compared to the parasite-free treatments (Fig. 2C, D). This equalising tendency might mean that the parasite makes demographic conditions in core and front populations more similar, and/or that potential general indirect effects of parasite-mediated selection partly override spatial selection on growth traits.

### Range expansion, but not parasitism influences swimming behaviour

Our data show that Paramecium can (evolve to) swim either straight and fast or more slowly and take more turns (Fig. 1). *Paramecium* from the range front treatment were clearly more of the slow type, consistent with previous observation [57]. This contradicts results in a very similar experiment with another ciliate [59] and observations in other biological systems, where populations at the range front show enhanced movement ability [34,37]. In our case, speed reduction was associated with increased swimming tortuosity. This latter tendency to change directions while swimming could represent an exploratory behaviour that might influence the probability of finding the dispersal corridors in our connected microcosms. While our range selection treatment had a strong impact on movement behaviour, there was no discernible effect of the parasite treatment. Thus, unlike for the other trait pairs, parasite-mediated selection does not seem to play a role in shaping the evolution of these traits. As stand-alone traits, speed and tortuosity are only weakly correlated with dispersal (Table S9), suggesting that the effect of parasite treatment on dispersal must operate through other aspects of movement.

### From eco to evo and back: Is the evolution with a parasite slowing down range expansions?

The evolution of dispersal syndromes typically produces ecological feedbacks that speed up range expansions [4]. Parasites, in contrast, may keep in check host populations and thereby limit or slow down range fronts, as generically shown for interspecific (antagonistic) interactions [5,11,80]. Parasites may indeed modify ecological dynamics by affecting host demography and dispersal during a range expansion, and consequently alter the strength or sign selection on these traits. Moreover, parasite resistance to is yet another trait under selection. The main purpose of this study was to investigate such eco-to-evo feedbacks and assess the possible impact of a parasite on the evolutionary trajectories of dispersal syndromes. The open question is how these evolutionary changes in turn feedback on the ecological dynamics of a range expansion, regarding the demography, dispersal or expansion speed. One of our main findings is that the presence of the parasite in the range front treatment favours higher resistance, but also leads to lower dispersal, when compared to the parasite-free treatment (Fig. 2A, B). Thus, an intuitive prediction is that evolution with the parasite slows down range expansion speed, due to this resistance - dispersal trade-off. To apply this prediction to natural systems is difficult because these might rarely be replicated with and without parasites. However, as already discussed above for the cane toad example, there are cases where resistance evolution does occur at the range expansion front [41]. And even though it is unknown how the resistance evolution in the toad is correlated with dispersal traits, it seems obvious that, in the face of a detrimental parasite (a lung worm), having the resistance will benefit the range expansion more than not having it [12], at least as long as parasites equally infect high- and low-dispersal individuals, preventing escape into enemy-free space.

Beyond these simple considerations, however, answers may be more complex [81], if we add a multi-trait perspective. As we have shown here, parasite-mediated selection may affect some dispersal syndrome components, but not others, or even modify the correlation between these components. Moreover, a full picture would require measurements of growth, dispersal, and movement in the presence of infection, thereby integrating aspects or virulence / tolerance or context-dependent dispersal [53,54]. In a next step towards a more comprehensive understanding of eco-evo feedbacks, future experiments can use multi-microcosm landscapes to actually measure range expansion speeds for the different types of evolved hosts, with and without parasite. Ideally, this would be accompanied by theory exploring the simultaneous evolution of dispersal and interaction traits [82] or accounting for additional factors, such as parasite evolution [53], dispersal plasticity [45,54] or the general upregulation of immune responses, known to occur during range expansion [76,77].

## Conclusion

Our selection experiment illustrates the possible impact of biotic interactions on evolutionary trajectories during range expansions. Parasite-mediated selection changed the structure of dispersal syndromes, shaping the phenotypic divergence between front and core populations for multiple traits, including dispersal, life-history and resistance. Here we focused on the single-sided host evolution, but dispersal syndromes also evolved in the parasite [53]. It could mean that our observed host changes may have an additional coevolutionary component. This may further complexify the interplay between spatial selection and biotic (co-)evolution, and make it even more difficult to predict the accompanying ecological feedbacks on range expansion dynamics or disease spread. Replicated range expansions under laboratory conditions represent one tool to disentangle these processes. Although being simplified abstractions of the real world, such experiments may also provide baseline information for applied issues of biocontrol or the monitoring of emerging diseases.

## Ethics

Not applicable

## Data accessibility

The data from this study are available at https://doi.org/10.5281/zenodo.7123494

## Authors’ contributions

OK, LN and GZ conceived the study. GZ, LN, MH, EAF, and OK designed the experiments. GZ, LN, CGB and OK performed the experimental work. GZ, LN, OK and EAF performed the statistical analysis. All authors interpreted the results. GZ and OK wrote the first draft of the manuscript and all authors commented on the final version.

## Acknowledgements

This is publication ISEM-YYYY-XXX of the Institut des Sciences de l’Evolution - Montpellier.

## Supplementary Information

### S1 Dispersal systems

We first filled the two-patch system with 9.5 mL of fresh growth medium and we then closed the corridor with a clamp. Secondly, we filled to 13 mL one of the two tubes, the core patch, using the paramecia and medium from an experimental selection line. The second tube, the front patch, was filled to 13 mL with fresh growth medium, and thus resulted empty at this stage. Thirdly, we removed the clamp and opened the corridor allowing the paramecia to actively disperse and swim from core to front patch or to stay in the core. After three hours, we closed the corridor tube and we estimated the population density by taking a 200 μL sample from core and front patch and counting the number of individuals under a dissecting microscope.

**Figure S1.**
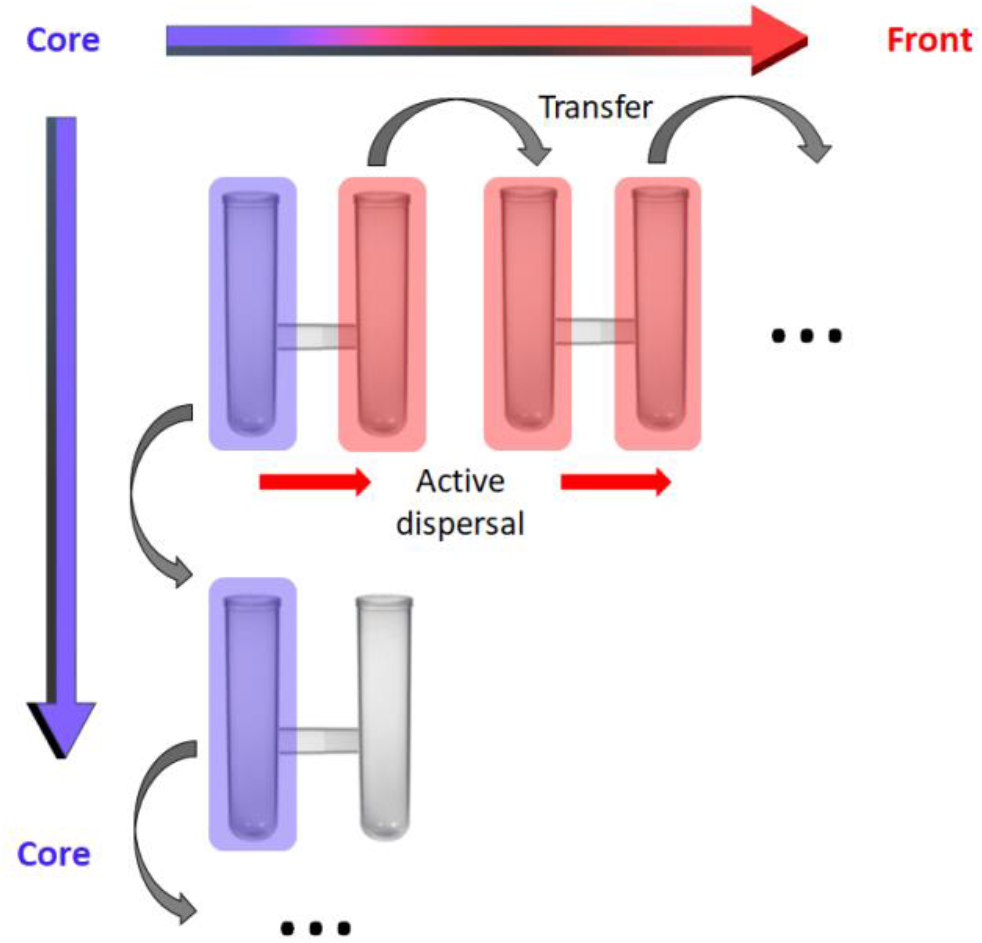
Dispersal arenas used for the range expansion dynamics with representation of how the front treatment was propagated. The arenas were composed from two patches (core and front) interconnected by 5-cm silicon tubing serving as a corridor. The *paramecia* could displace from core to front patch through active dispersal once we removed the clamp blocking the corridor. In the long-term experiment, we alternated short episodes of dispersal (3h) with periods of population growth and maintenance (1 week = 1 cycle). The same protocol was repeated for the core and front treatment with the parasite.

### S2 Extraction protocol

Inocula were prepared concentrating by centrifugation infected paramecia (35000 RPM for 20 minutes) in Falcon tubes filled with 15 mL of medium. After removing the supernatant, the concentrated individuals were transferred into 1.5 mL Eppendorf tubes containing 1 mm glass beads, and they were vortexed and crushed using a Qiagen TissuLyser (1.45 minutes at 30 oscillation frequency). The released infectious forms were then counted using a hemocytometer at 200 x magnification under a microscope (Leica DM LB2), and their concentration was adjusted with sterile water.

### S3 Design phenotypic trait assays

**Figure S3.**
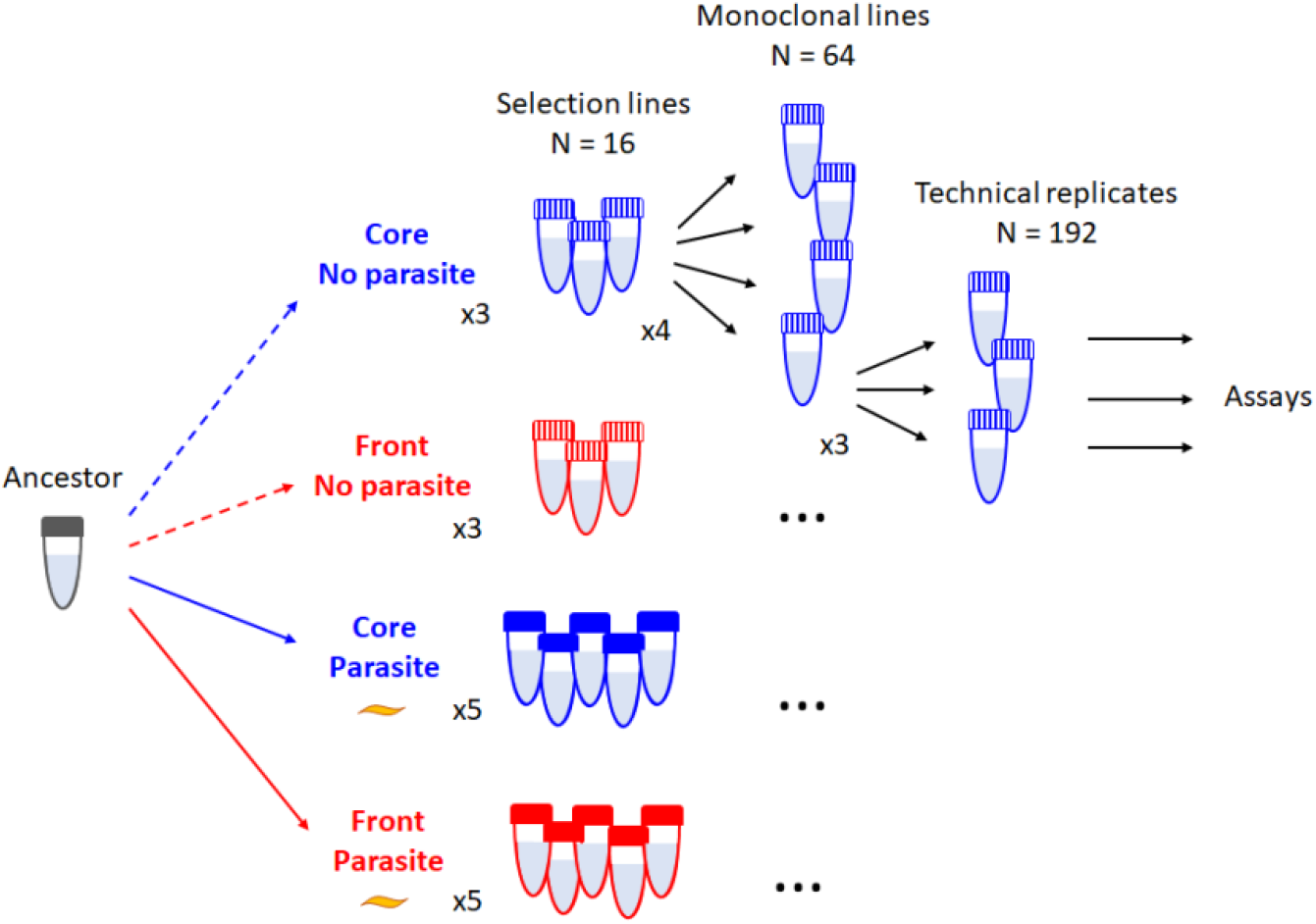
Experimental design of the phenotypic trait assays (16 × 4 × 3 tubes). Some technical replicates were lost during the manipulation, and the final number was 180. This corresponded to the total of 63 monoclonal lines that were used in the analysis.

### S4 Long-Term cycles

**Figure S4.**
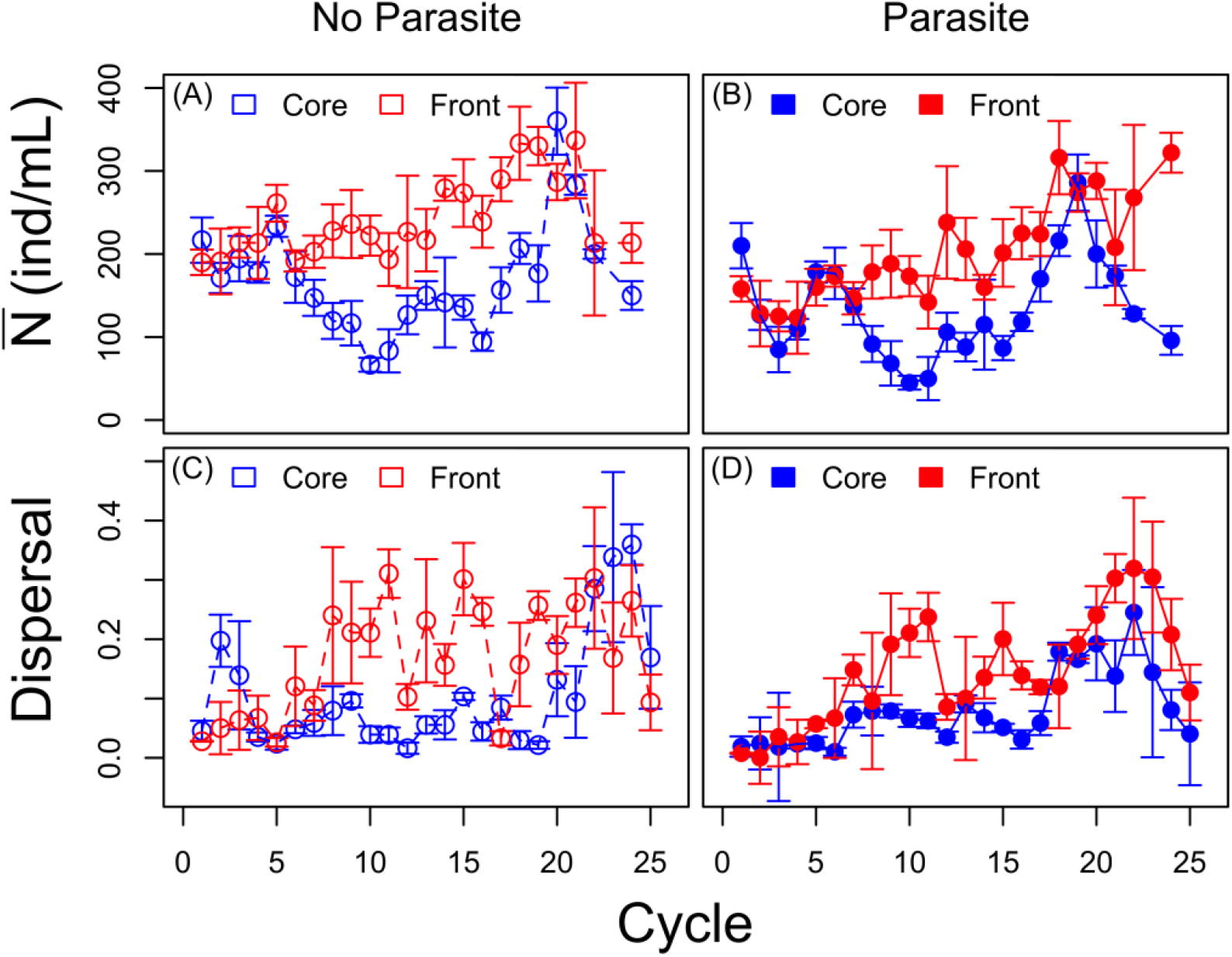
Density (individuals per mL) and dispersal (proportion of individuals dispersing in 3 hours) dynamics during the first 25 cycles of the long-term range expansion experiment for (A)-(C) populations not infected (empty symbols), and (B)-(D) infected (full symbols). Each point is the selection lines mean with the standard error, core treatment is in blue and front treatment in red.

### S5 Dispersal rate

We followed a similar procedure to what was previously described for the dispersal systems. In short, we filled the 3-patch systems with 40 mL of fresh growth medium, we closed the two corridors with clamps, and then added the entire 25 mL of the final tube to the central compartment. Left and right tubes were filled with fresh growth medium to have the same amount of volume of the central tube, then the corridors were opened. We allowed the paramecia to actively disperse in both directions for three hours before closing the dispersal corridors again. We sampled 500 μL from the core patch, 3 mL from both front patches and we estimated population densities.

**Figure S5.**
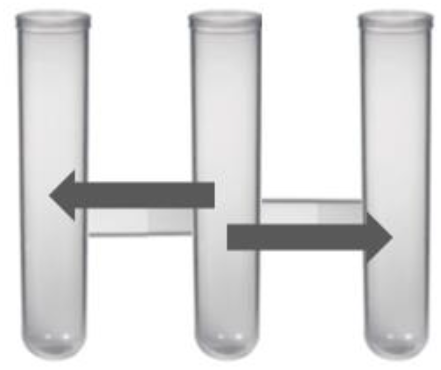
Linear 3-patch arena used to measure dispersal rate after evolution (3 interconnected 50 mL Falcon tubes). The Paramecium dispersed from the middle to the two outer tubes (arrows). Similarly to the protocol used during the long-term experiment, we opened the corridors for 3 h. Dispersal rates were estimated by sampling and counting the Paramecium from the central tube (500 μl) and from the combined two outer tubes (3 mL). We did not control for the density of Paramecium placed in the middle tube of the dispersal arena. However, preliminary analysis showed no significant effect of density on dispersal rate (F_1,54_= 0.69, n.s.), and the covariate was therefore omitted from further analyses.

## Tables

**Table S1.** PCA Eigenvalues and variance explained. PCA based on values for 6 traits measured for 63 monoclonal lines from 4 long-term treatments (for details, see main text).

|  | PC1 | PC2 | PC3 | PC4 | PC5 | PC6 |
| --- | --- | --- | --- | --- | --- | --- |
| Eigenvalues | 2.0876 | 1.3090 | 1.0711 | 0.6416 | 0.5950 | 0.2964 |
| Proportion variance explained | 0.3479 | 0.2182 | 0.1785 | 0.1070 | 0.0991 | 0.0492 |
| Cumulative proportion | 0.3479 | 0.5661 | 0.7446 | 0.8516 | 0.9507 | 1.0000 |

**Table S2.**
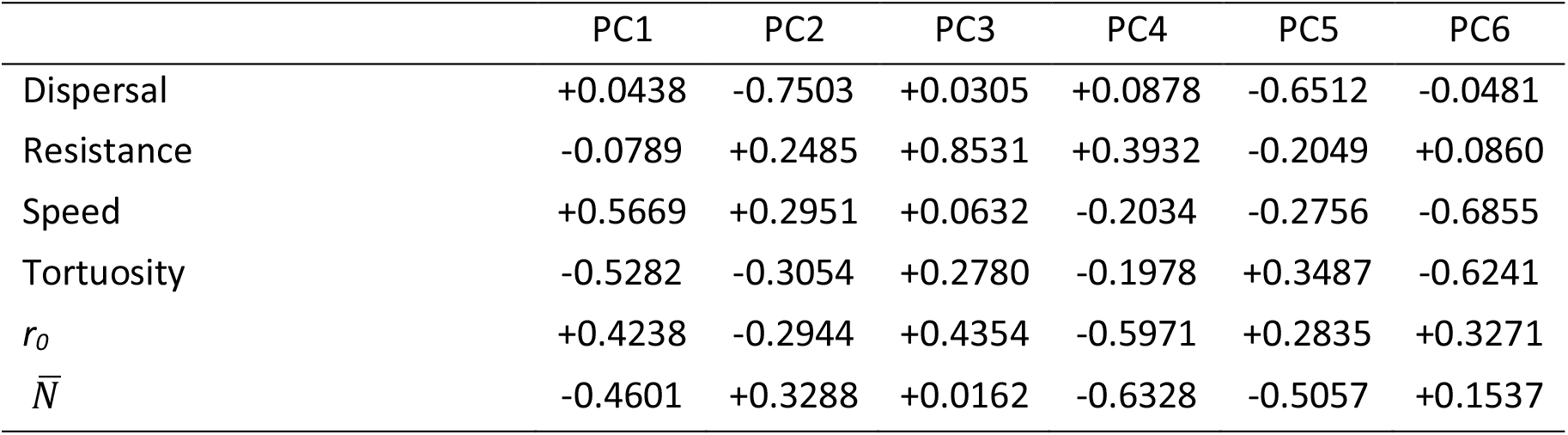
Loading values of the PCA for 6 host traits.

**Table S3.** Single trait analysis of dispersal, testing effects of range expansion (front vs core) and parasitism (absence / presence of parasite) treatments. Rows are the different models; the best model is highlighted in bold. Columns are the explanatory variables included with the corresponding WAIC, standard error of the WAIC and WAIC weights for each model. The RI row indicates the relative importance of the explanatory variables.

|  | Range expansion | Parasitism | Range * Parasitism | WAIC | SE | WAIC weight |
| --- | --- | --- | --- | --- | --- | --- |
| Model 1 |  |  |  | 155.6 | 11.9 | 0.14 |
| **Model 2** |  | **X** |  | **154.2** | **12.7** | **0.28** |
| Model 3 | X |  |  | 156.0 | 11.9 | 0.11 |
| Model 4 | X | X |  | 154.8 | 12.6 | 0.21 |
| Model 5 | X | X | X | 154.3 | 12.7 | 0.26 |
| RI | 0.58 | 0.75 | 0.26 |  |  |  |

**Table S4.**
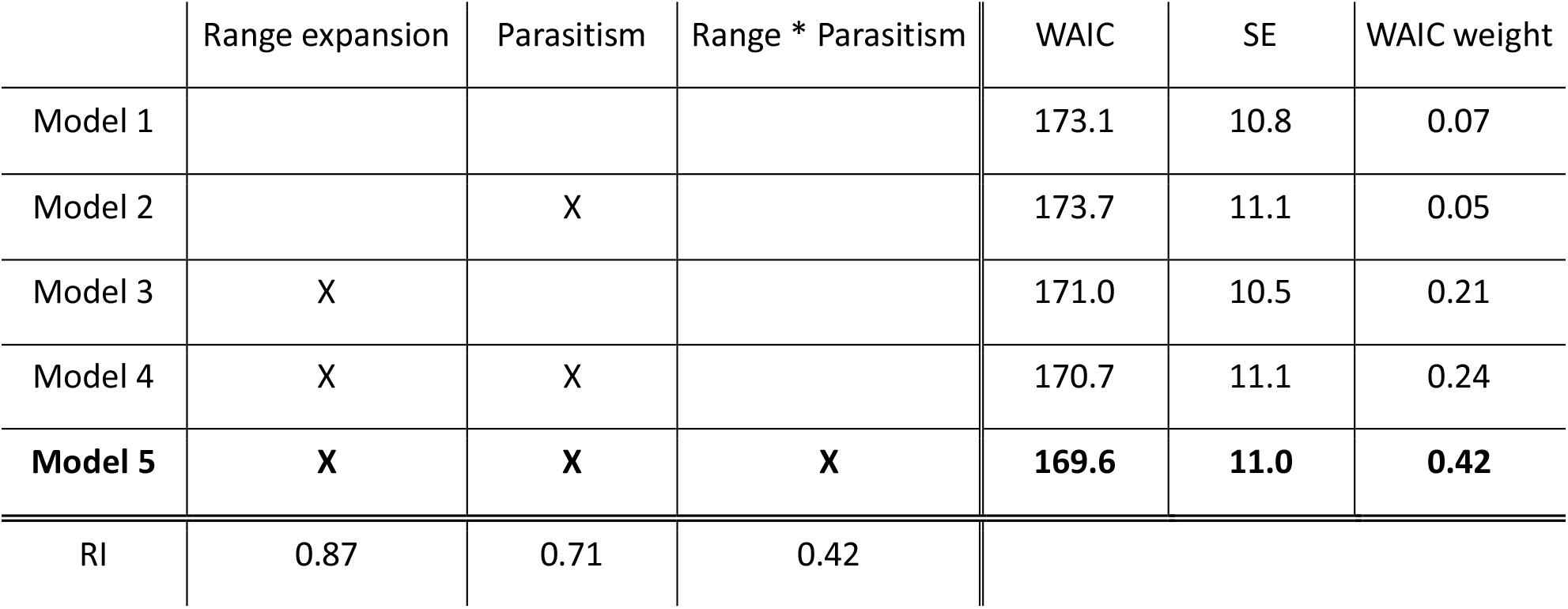
Single trait analysis of resistance, testing effects of range expansion (front vs core) and parasitism (absence / presence of parasite) treatments. Rows are the different models; the best model is highlighted in bold. Columns are the explanatory variables included with the corresponding WAIC, standard error of the WAIC and WAIC weights for each model. The RI row indicates the relative importance of the explanatory variables.

**Table S5.** Single trait analysis of *r*_*0*_, testing effects of range expansion (front vs core) and parasitism (absence / presence of parasite) treatments. Rows are the different models; the best model is highlighted in bold. Columns are the explanatory variables included with the corresponding WAIC, standard error of the WAIC and WAIC weights for each model. The RI row indicates the relative importance of the explanatory variables.

|  | Range expansion | Parasitism | Range * Parasitism | WAIC | SE | WAIC weight |
| --- | --- | --- | --- | --- | --- | --- |
| Model 1 |  |  |  | 181.7 | 20.1 | 0.17 |
| Model 2 |  | X |  | 182.3 | 19.8 | 0.12 |
| Model 3 | X |  |  | 180.8 | 19.7 | 0.26 |
| Model 4 | X | X |  | 181.5 | 19.5 | 0.18 |
| **Model 5** | **X** | **X** | **X** | **180.8** | **19.2** | **0.26** |
| RI | 0.70 | 0.56 | 0.26 |  |  |  |

**Table S6.** Single trait analysis of 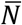, testing effects of range expansion (front vs core) and parasitism (absence / presence of parasite) treatments. Rows are the different models; the best model is highlighted in bold. Columns are the explanatory variables included with the corresponding WAIC, standard error of the WAIC and WAIC weights for each model. The RI row indicates the relative importance of the explanatory variables.

|  | Range expansion | Parasitism | Range * Parasitism | WAIC | SE | WAIC weight |
| --- | --- | --- | --- | --- | --- | --- |
| Model 1 |  |  |  | 98.4 | 14.2 | 0.18 |
| Model 2 |  | X |  | 98.1 | 13.3 | 0.21 |
| Model 3 | X |  |  | 98.5 | 14.4 | 0.17 |
| **Model 4** | **X** | **X** |  | **97.9** | **13.4** | **0.23** |
| Model 5 | X | X | X | 98.0 | 13.8 | 0.22 |
| RI | 0.62 | 0.66 | 0.22 |  |  |  |

**Table S7.** Single trait analysis of swimming speed, testing effects of range expansion (front vs core) and parasitism (absence / presence of parasite) treatments. Rows are the different models; the best model is highlighted in bold. Columns are the explanatory variables included with the corresponding WAIC, standard error of the WAIC and WAIC weights for each model. The RI row indicates the relative importance of the explanatory variables.

|  | Range expansion | Parasitism | Range * Parasitism | WAIC | SE | WAIC weight |
| --- | --- | --- | --- | --- | --- | --- |
| Model 1 |  |  |  | 182.4 | 19.3 | 0.04 |
| Model 2 |  | X |  | 184.0 | 19.5 | 0.02 |
| **Model 3** | **X** |  |  | **176.7** | **18.5** | **0.62** |
| Model 4 | X | X |  | 178.8 | 18.8 | 0.22 |
| Model 5 | X | X | X | 180.4 | 18.5 | 0.10 |
| RI | 0.94 | 0.34 | 0.10 |  |  |  |

**Table S8.** Single trait analysis of swimming tortuosity, testing effects of range expansion (front vs core) and parasitism (absence / presence of parasite) treatments. Rows are the different models; the best model is highlighted in bold. Columns are the explanatory variables included with the corresponding WAIC, standard error of the WAIC and WAIC weights for each model. The RI row indicates the relative importance of the explanatory variables.

|  | Range expansion | Parasitism | Range * Parasitism | WAIC | SE | WAIC weight |
| --- | --- | --- | --- | --- | --- | --- |
| Model 1 |  |  |  | 181.6 | 11.0 | 0.03 |
| Model 2 |  | X |  | 182.1 | 11.0 | 0.02 |
| **Model 3** | **X** |  |  | **176.5** | **11.3** | **0.39** |
| Model 4 | X | X |  | 176.8 | 10.9 | 0.34 |
| Model 5 | X | X | X | 177.7 | 10.8 | 0.22 |
| RI | 0.95 | 0.56 | 0.22 |  |  |  |

**Table S9.** Mean correlation coefficients for selected pairs of host traits (± 95% compatibility intervals obtained from posterior distributions).

| Pair of traits | Range expansion | Parasitism | Mean correlation coefficient ± 95% CI |
| --- | --- | --- | --- |
| Dispersal - Resistance | Core | No | +0.155 (-0.246; +0.515) |
| Dispersal - Resistance | Core | Yes | -0.091 (-0.459; +0.299) |
| Dispersal - Resistance | Front | No | -0.387 (-0.761; +0.168) |
| Dispersal - Resistance | Front | Yes | -0.440 (-0.648; - 0.183) |
| $r_0 - \bar{N}$ | Core | No | +0.264 (-0.336; +0.725) |
| $r_0 - \bar{N}$ | Core | Yes | -0.459 (-0.774; -0.021) |
| $r_0 - \bar{N}$ | Front | No | +0.080 (-0.519; 0.643) |
| $r_0 - \bar{N}$ | Front | Yes | -0.702 (-0.888; -0.399) |
| Speed - Tortuosity | Core | No | -0.915 (-0.978; -0.680) |
| Speed - Tortuosity | Core | Yes | -0.409 (-0.738; +0.067) |
| Speed - Tortuosity | Front | No | -0.319 (-0.777; +0.315) |
| Speed - Tortuosity | Front | Yes | -0.841 (-0.944; -0.652) |
| Dispersal - Tortuosity | Core | No | +0.043 (-0.419; +0.506) |
| Dispersal - Tortuosity | Core | Yes | +0.246 (-0.210; +0.636) |
| Dispersal - Tortuosity | Front | No | +0.302 (-0.253; +0.726) |
| Dispersal - Tortuosity | Front | Yes | +0.206 (-0.119; +0.489) |
| Dispersal - Speed | Core | No | +0.206 (-0.119; +0.489) |
| Dispersal - Speed | Core | Yes | -0.136 (-0.569; +0.329) |
| Dispersal - Speed | Front | No | -0.249 (-0.691; +0.319) |
| Dispersal - Speed | Front | Yes | -0.248 (-0.528; +0.067) |
| Resistance- $r_0$ | Core | No | +0.363 (-0.087; +0.694) |
| Resistance- $r_0$ | Core | Yes | +0.330 (-0.057; +0.635) |
| Resistance- $r_0$ | Front | No | -0.307 (-0.763; +0.297) |
| Resistance- $r_0$ | Front | Yes | -0.135 (-0.441; +0.200) |

